# Modeling the antigenic evolution of seasonal influenza viruses with PREDAC for vaccine strain selection in the season of 2025

**DOI:** 10.1101/2024.09.08.611938

**Authors:** Yousong Peng, Xiangjun Du, Liu Mi, Xiao Ding, Jing Meng, Wenjie Han, Lei Yang, Weijuan Huang, Taijiao Jiang, Dayan Wang

## Abstract

**Introduction:** The antigenic evolution of seasonal influenza viruses have been reported to be discontinuous as clusters. Timely identification of antigenic clusters could help for influenza virus vaccine strain selections.

**Methods:** Here, we revealed the antigenic evolution of seasonal influenza viruses, including A(H3N2), A(H1N1pdm) and B-Victoria, using our previously developed PREDAC methods, and identified antigenic clusters of seasonal influenza viruses based on existing HA protein sequences in GISAID database.

**Results:** During the last six months (2024.2∼2024.7), for the influenza A(H3N2) virus, the antigenic clusters DA21 (A/Darwin/6/2021-like and A/Massachusetts/18/2022-like) continuously dominated and nearly all viruses are antigenic similar to the current vaccine strain A/Massachusetts/18/2022; for the influenza A(H1N1pdm) virus, the antigenic cluster SY21 (A/Sydney/5/2021-like and A/Victoria/4897/2022-like) dominated and nearly all viruses are antigenic similar to the current vaccine strain A/Victoria/4897/2022; for the influenza B-Victoria, antigenic clusters AU21 (B/Austria/1359417/2021-like) dominated and nearly all viruses are antigenic similar to the current vaccine strain B/Austria/1359417/2021.

**Conclusion:** The analysis suggests that there is no need to update the vaccine strains in the flu season of 2025 on the aspect of antigenicity.

## Introduction

Vaccination is currently the most effective way to fight against the influenza viruses. Unfortunately, the influenza virus frequently changes its antigen by rapid mutations, which leads to decreased efficiency or even failure of vaccines. To improve the vaccine efficiency, it is necessary to surveillance the antigenic variation of the virus and update vaccine strains when significant antigenic variation happens. We previously developed a computational method PREDAC (PREDict Antigenic Cluster) for predicting antigenic clusters, i.e., a group of viruses with similar antigenicity, of influenza viruses based on HA protein sequences ^1-8^. PREDAC not only achieves a high accuracy in the prediction of antigenic variations under the machine learning algorithm but also models the dynamics of antigenic patterns of influenza viruses in a form of antigenic clusters (a group of viruses with similar antigenicity). Based on PREDAC, we further proposed a novel framework for flu vaccine strain recommendations^1^. We showed that coupling PREDAC with large-scale HA sequencing of influenza A(H3N2) viruses in the mainland China could help improve the vaccine recommendation of the virus. The PREDAC method has been used for vaccine recommendations of influenza viruses since 2013. Here, we revealed the antigenic evolution of seasonal influenza viruses, including A(H3N2), A(H1N1pdm) and B-Victoria, using the PREDAC methods, and identified antigenic clusters of seasonal influenza viruses based on existing HA protein sequences. It would provide insights into the influenza virus vaccine recommendation in the coming flu season.

## Materials and Methods

### Data

The HA protein sequences of the influenza A(H1N1pdm), A(H3N2) and the Victoria clade of the influenza B virus were downloaded from GISAID database (https://gisaid.org/) as of July 31^th^, 2024.

### Antigenic modeling and analysis

The antigenic clustering of HA protein sequences was achieved with the PREDAC series methods^1,2^. Specifically, for the influenza A(H3N2) virus, the antigenic clustering results were directly obtained from the web-server of PREDAC-H3^5^; for the influenza A(H1N1pdm) virus, the method of antigenic clustering was adapted from Liu’s work^8^; for the Victoria clade of the influenza B virus, the method of antigenic clustering was newly developed.

### Phylogenetic analysis

The extensive set of hemagglutinin (HA) nucleotide genomic sequences were refined to enhance the clarity of the phylogenetic tree, employing the following criteria: i) all sequences collected in 2024 were included; ii) for sequences sampled before 2024 with distinct combinations of antigenic cluster, collection country, and collection date, non-redundant sequences were derived using cd-hit with a sequence similarity cutoff of 0.986. Phylogenetic trees were constructed using FastTreeDBI^9^ with 100 bootstraps (https://www.phylo.org/index.php/tools/fasttree_xsede.html). The Maximum Likelihood model was employed for building the phylogenetic tree with nucleotide sequences. The resultant trees were visualized using the R packages ggtree, ggplot2, and ape.

### Visualization

The bar plots and scatter plots were drawn in R (v4.0.3). The antigenic correlation network of influenza viruses was visualized with Cytoscape (v3.8.0)^10^.

## Results

### Modeling the antigenic evolution of seasonal influenza A(H3N2) viruses

#### Location and time distribution of HA protein sequences of influenza A(H3N2) viruses since 2022

The monthly number of HA protein sequences of influenza A(H3N2) viruses from 2022 was shown in Figure S1 by region. It can roughly reflect the epidemic of the virus in each region. During the 2024 season, most HA protein sequences were from Europe and North America.

#### Antigenic clustering of influenza A(H3N2) viruses

All the HA protein sequences of influenza A(H3N2) viruses were antigenically clustered with the PREDAC-like method. As shown in Figure 1, the antigenic cluster DA21(A/Darwin/6/2021-like) dominated in the last three seasons. The recommended vaccine strains in recent two flu seasons, i.e., A/Thailand/8/2022(egg) and A/Massachusetts/18/2022(cell), were also classified to be DA21. The clade composition of viruses isolated in recent two years in the DA21 was analyzed. As shown in Figure 1C, most viruses (82.3%) belonged to the clade 3C.2a1b.2a.2a.3a.1, which was the predominant clade in the last flu season.

**Figure 1.**
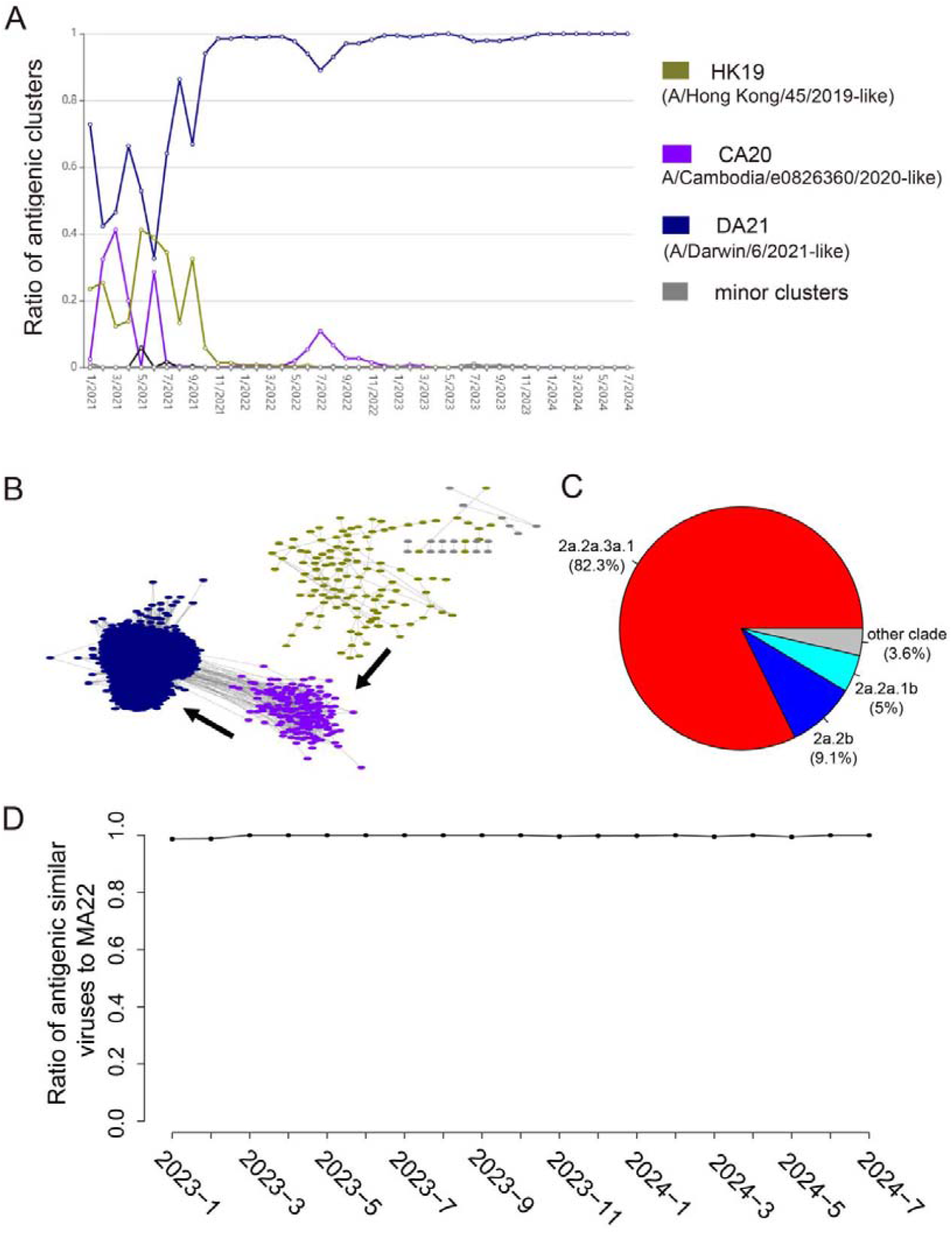
The antigenic clustering of influenza A(H3N2) viruses. (A) Monthly dynamics of antigenic clusters in the periods of 2021-2024. The colored lines refer to the ratio of antigenic clusters, while the gray lines refer to the minor clusters. (B) The antigenic correlation network of viruses in the last three years (2021-2024). Each node referred to a virus isolate. The nodes were colored by antigenic clusters. The antigenic clusters were named by the vaccine strain according to the figure legend. For clarity, only 4% of the non-redundant sequences were randomly selected and shown in the network. (C) The clade composition of viruses isolated in recent two years in the DA21. (D) The monthly ratio of antigenic similar viruses to the latest vaccine strain A/Massachusetts/18/2022 in the last two years.

#### Phylogenetic analysis of recent influenza A(H3N2) viruses

The phylogenetic tree for the HA genomic sequences of representative influenza A(H3N2) viruses collected since 2021 was constructed. Strains were color-coded according to their antigenic clusters (see the figure legend). As depicted in Figure 2, viruses within the clade 3C.2a1b.2a.2a.3a.1 were categorized under the antigenic cluster DA21. Notably, nearly all viruses collected in the last half year were assigned to the DA21 cluster (see right side of the figure).

**Figure 2.**
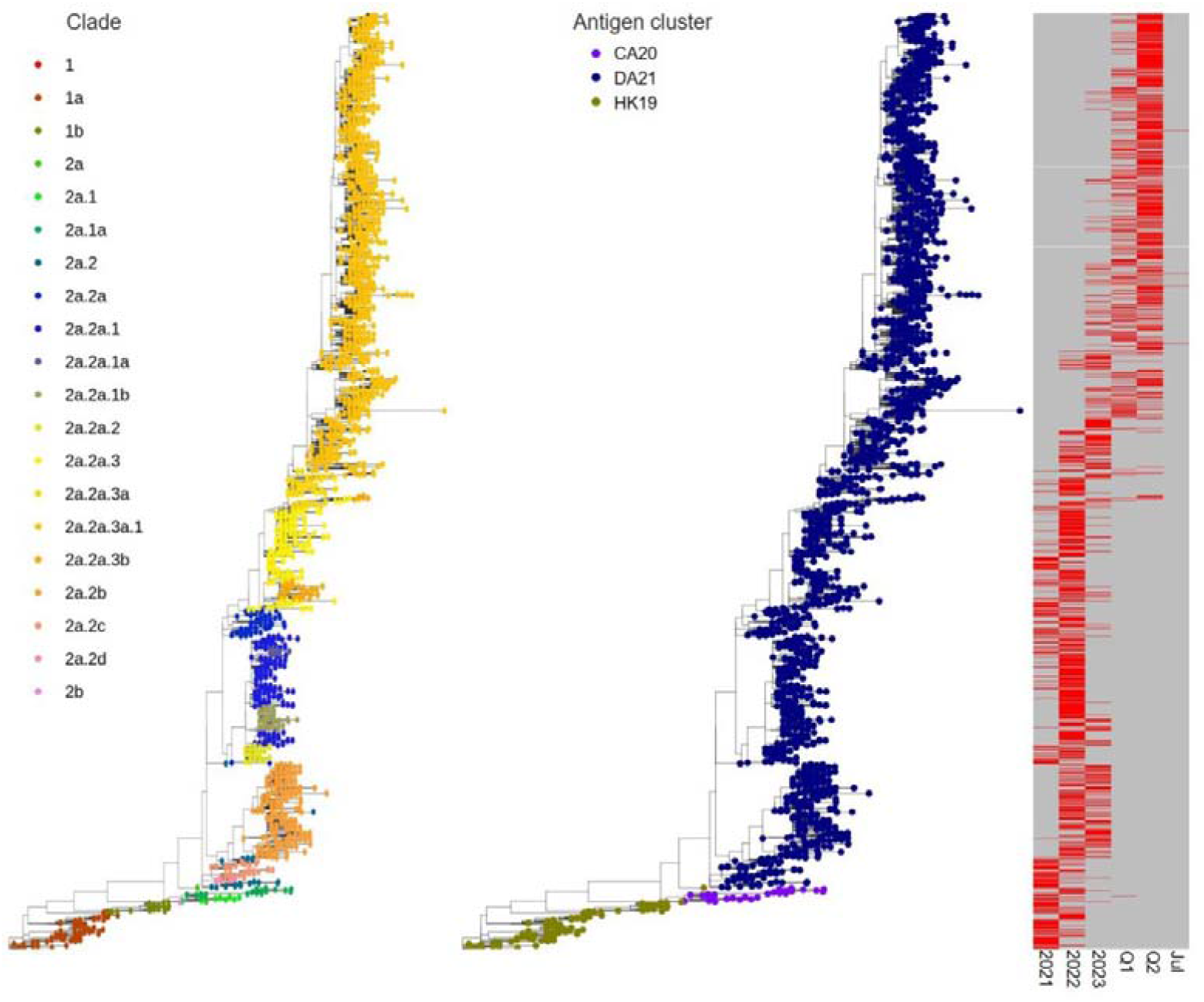
The phylogenetic tree represents recent influenza A(H3N2) viruses, with colors indicating the clade (left figure) or antigenic clusters (middle figure) as per the legend in the top-left corner of the figure. The right figure refers to the isolation time of viruses in the phylogenetic tree. Q1 and Q2 refer to the first and second quarter of this year. Clades were determined using the Nextclade pipeline.

#### Monthly ratio of antigenic similar viruses to the latest vaccine strain

Then,we analyzed the monthly ratio of antigenic similar viruses to the latest vaccine strain A/Massachusetts/18/2022 in the last two years. As shown in Figure 1D, nearly 100% of viruses were antigenic similar to the current vaccine strain, suggesting that the current vaccine strain may protect well against viruses in the last two years.

### Modeling the antigenic evolution of seasonal influenza A(H1N1pdm) viruses

#### Location and time distribution of HA protein sequences of influenza A(H1N1pdm) viruses since 2021

The monthly number of HA protein sequences of influenza A(H1N1pdm) viruses since 2021 was shown in Figure S2 by region. It can roughly reflect the epidemic of the virus in each region. During the 2024 season, most HA protein sequences were from Europe and North America.

#### Antigenic clustering of influenza A(H1N1) viruses

All the HA protein sequences of influenza A(H1N1pdm) viruses were antigenically clustered with the PREDAC-like method. As shown in Figure 3A&B, the antigenic cluster SY21(A/Sydney/5/2021-like) dominated in the last three seasons. The recommended vaccine strains in recent two flu seasons, i.e., A/Victoria/4897/2022(egg) and A/Wisconsin/67/2022(cell), were also classified to be SY21. The clade composition of viruses isolated in recent two years in the SY21 was analyzed. As shown in Figure 3C, nearly all viruses belonged to the clade 6B.1A.5a.2a (65.1%) and its sub-clade 6B.1A.5a.2a.1 (34.5%), which were the predominant clades in the last flu season.

**Figure 3.**
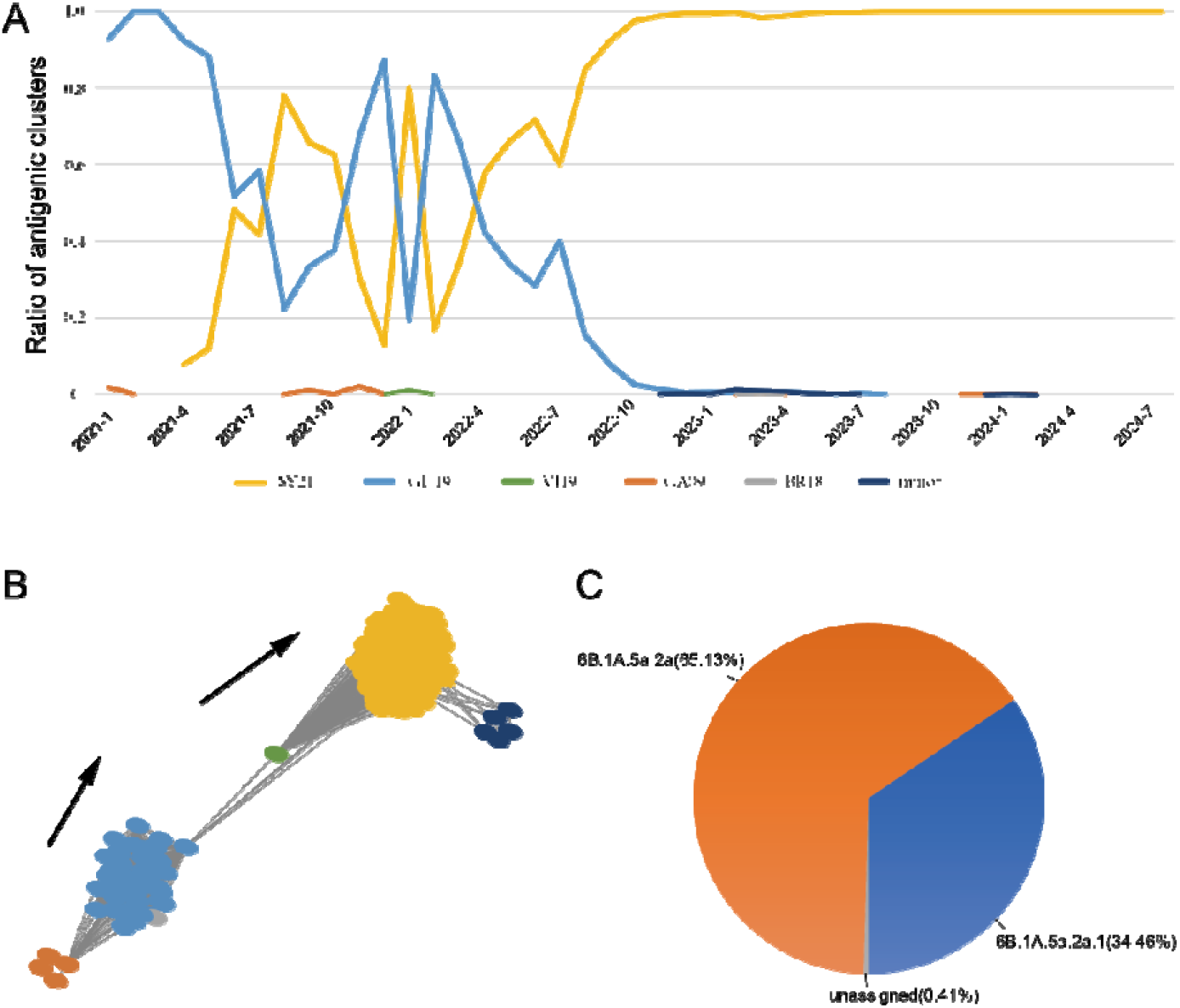
The antigenic clustering of influenza A(H1N1pdm) viruses. (A) Monthly dynamics of antigenic clusters in the periods of 2021-2024. The colored lines refer to the ratio of antigenic clusters, while the dark blue lines refer to the minor clusters. (B) The antigenic correlation network of viruses in the last three years (2021-2024). Each node referred to a virus isolate. The nodes were colored by antigenic clusters. The antigenic clusters were named by the vaccine strain according to the figure legend. For clarity, only 4% of the non-redundant sequences were randomly selected and shown in the network. (C) The clade composition of viruses isolated in recent two years in the SY21.

#### Phylogenetic analysis of recent influenza A(H1N1pdm) viruses

The phylogenetic tree for the HA genomic sequences of representative influenza A(H1N1pdm) viruses collected since 2021 was constructed. Strains were color-coded according to their antigenic clusters. As depicted in Figure S3, viruses within the clade 6B.1A.5a.2a and its sub-clade 6B.1A.5a.2a.1 were categorized under the antigenic cluster SY21. Notably, a majority of viruses collected in the last half year were associated with the SY21 cluster.

#### Monthly ratio of antigenic similar viruses to the latest vaccine strain

Then, we analyzed the monthly ratio of antigenic similar viruses to the latest vaccine strain A/Victoria/4897/2022 in the last two years. As shown in Figure S4, nearly 100% of viruses were antigenic similar to the current vaccine strain, suggesting that the current vaccine strain may protect well against viruses in the last two years.

### Modeling the antigenic evolution of seasonal influenza B (Victoria clade) viruses *Location and time distribution of HA protein sequences of influenza B (Victoria clade) viruses since 2022*

The monthly number of HA protein sequences of influenza B (Victoria clade) viruses since 2022 was shown in Figure S5 by region. It can roughly reflect the epidemic of the virus in each region. During the 2024 season, most HA protein sequences were from North America and Europe.

### Antigenic clustering of influenza B(Victoria clade) viruses

All the HA protein sequences of influenza B(Victoria clade) viruses were antigenically clustered with the PREDAC-like method. As shown in Figure 4, the antigenic cluster AU21(B/Austria/1359417/2021-like) dominated in the last three seasons. The clade composition of viruses isolated in recent two years in the AU21 was analyzed. All viruses (100%) belonged to the clade V1A.3a.2, which was the predominant clade in the last flu season.

**Figure 4.**
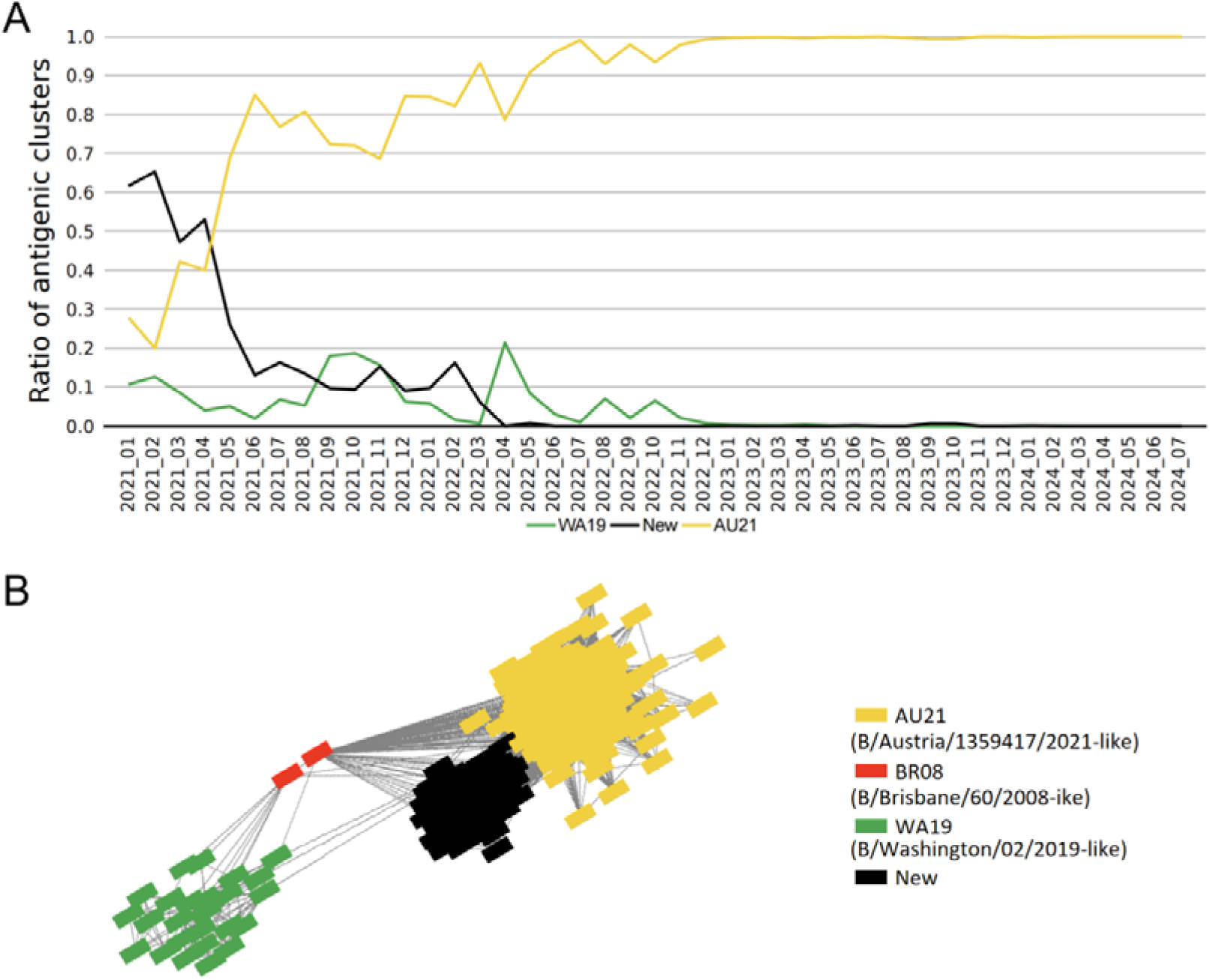
The antigenic clustering of influenza B (Victoria clade) viruses. (A) The monthly dynamics of antigenic clusters of influenza B(Victoria clade) viruses in the periods of 2021-2024. For clarity, only the dominant antigenic clusters were shown here in colored lines according to the figure legend in the bottom of the figure. (B) The antigenic correlation network of B (Victoria clade) viruses in recent three years. Each node referred to a virus isolate. The nodes were colored by antigenic clusters. The antigenic clusters were named by the vaccine strain according to the figure legend. For clarity, only the non-redundant sequences were selected and shown in the network.

### Phylogenetic analysis of recent influenza B(Victoria clade) viruses

The phylogenetic tree for the HA genomic sequences of representative influenza B (Victoria clade) viruses collected since 2023 was constructed. Strains were color-coded according to their antigenic clusters. As depicted in Figure S6, viruses within the clade V1A.3a.2 were categorized under the antigenic cluster AU21. Notably, all viruses collected in the last half year were associated with the AU21 cluster.

### Monthly ratio of antigenic similar viruses to the latest vaccine strain

Then,we analyzed the monthly ratio of antigenic similar viruses to the latest vaccine strain B/Austria/1359417/2021 in the last two years. As shown in Figure S7, nearly 100% of viruses were antigenic similar to the current vaccine strain, suggesting that the current vaccine strain may protect well against viruses in the last two years.

## Conclusion

During the last six months (2024.2∼2024.7), for all three type or subtypes of seasonal influenza viruses, viruses were antigenic similar to current vaccine strains, which suggests that it is not necessary to update the current vaccine composition on the aspect of antigenicity.

## Acknowledgments

We thank numerous submitters for the timely sharing of influenza virus sequence data in GISAID.

## Notes

### Competing Interest Statement

The authors have declared no competing interest.

