## Supplementary Materials for "Modeling the antigenic evolution of seasonal influenza viruses with PREDAC for vaccine strain selection in the season of 2025"

^1^Bioinformatics Center, College of Biology, Hunan Provincial Key Laboratory of Medical Virology, Hunan University, Changsha 410082, China; ^2^Department of Biostatistics and Systems Biology, School of Public Health (Shenzhen), Sun Yat-sen University, Shenzhen, Guangdong, 510275, China; ^3^Jiangsu Institute of Clinical Immunology, The First Affiliated Hospital of Soochow University, Suzhou, China; ^4^Institute of Systems Medicine, Chinese Academy of Medical Sciences & Peking Union Medical College, Beijing, China; ^5^Suzhou Institute of Systems Medicine, Suzhou, China; ^6^ National Institute for Viral Disease Control and Prevention, China CDC, Beijing 102206, China; ^7^Guangzhou National Laboratory, Guangzhou, China

**Figure S1**. The monthly number of HA protein sequences since 2022. E.SE.Asia includes countries in East and South-East Asia except China. Other.Asia refers to all countries in Asia except those in East and South-East Asia.

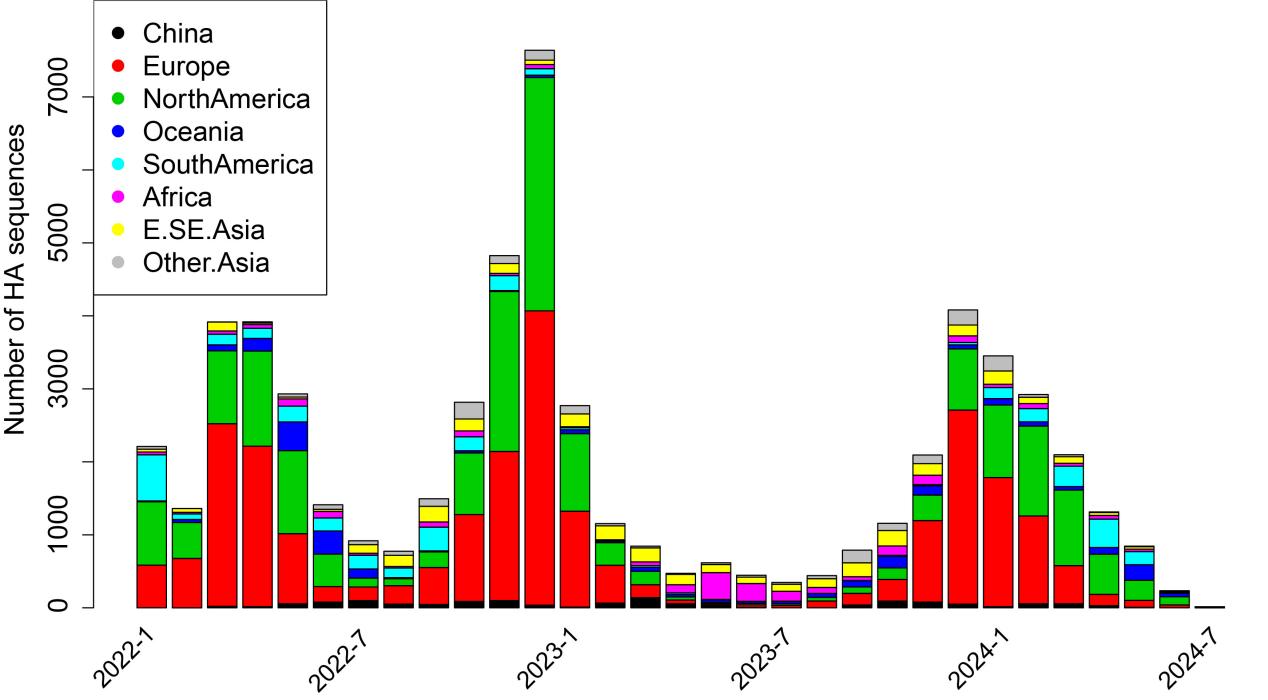

**Figure S2**. The monthly number of HA protein sequences since 2021. E.SE.Asia includes countries in East and South-East Asia except China. Other.Asia refers to all countries in Asia except those in East and South-East Asia.

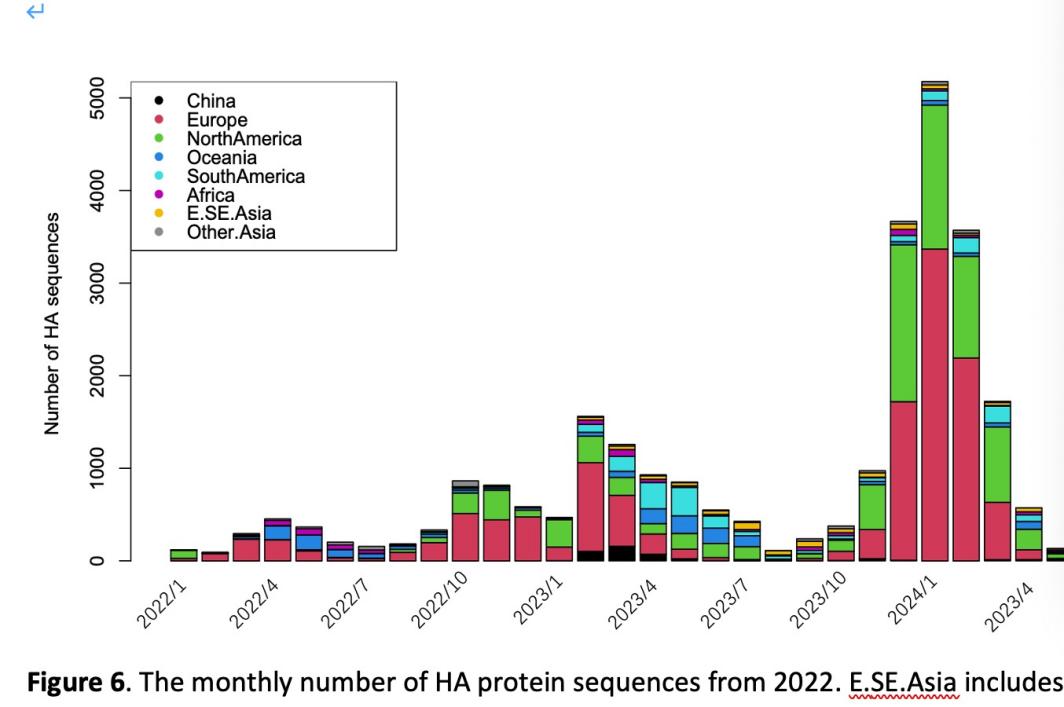

**Figure S3**. The phylogenetic tree represents recent influenza A(H1N1pdm) viruses, with colors indicating the clade (left figure) or antigenic clusters (middle figure) as per the legend in the top-left corner of the figure. The right figure refers to the isolation time of viruses in the phylogenetic tree. Q1 and Q2 refer to the first and second quarter of this year. Clades were determined using the Nextclade pipeline.

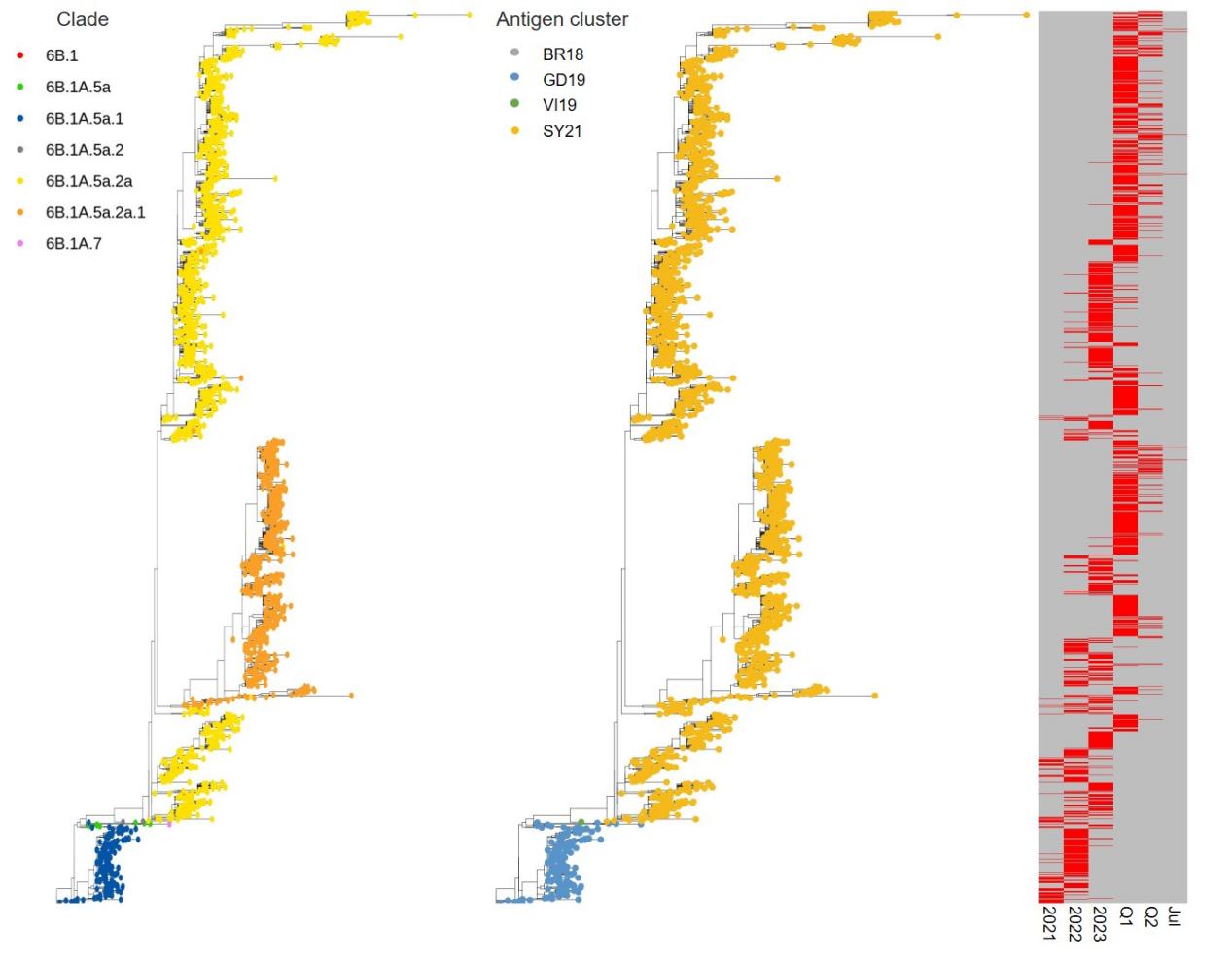

**Figure S4**. The monthly ratio of antigenic similar viruses to the latest vaccine strain A/Victoria/4897/2022 in the last two years.

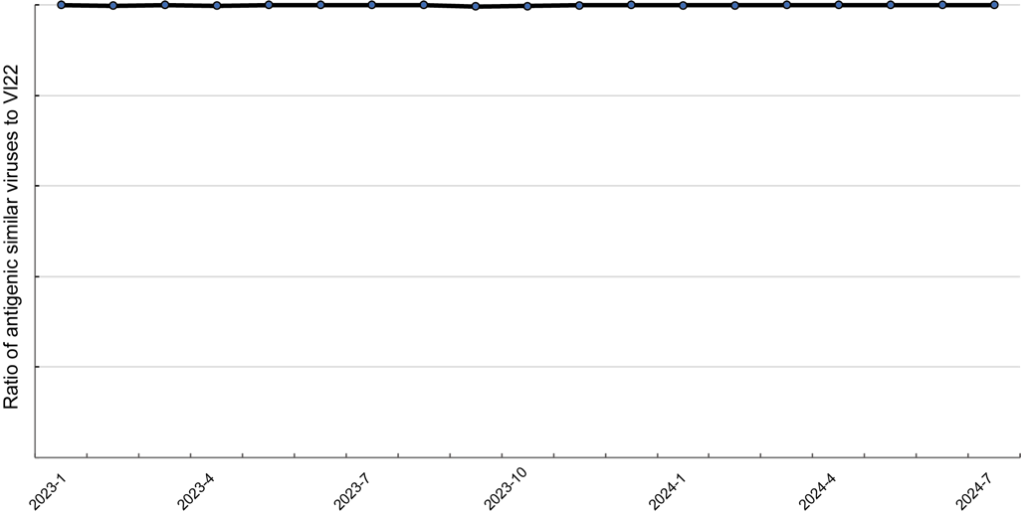

**Figure S5**. The monthly number of HA protein sequences since 2022. E.SE.Asia includes countries in East and South-East Asia except China. Other.Asia refers to all countries in Asia except those in East and South-East Asia.

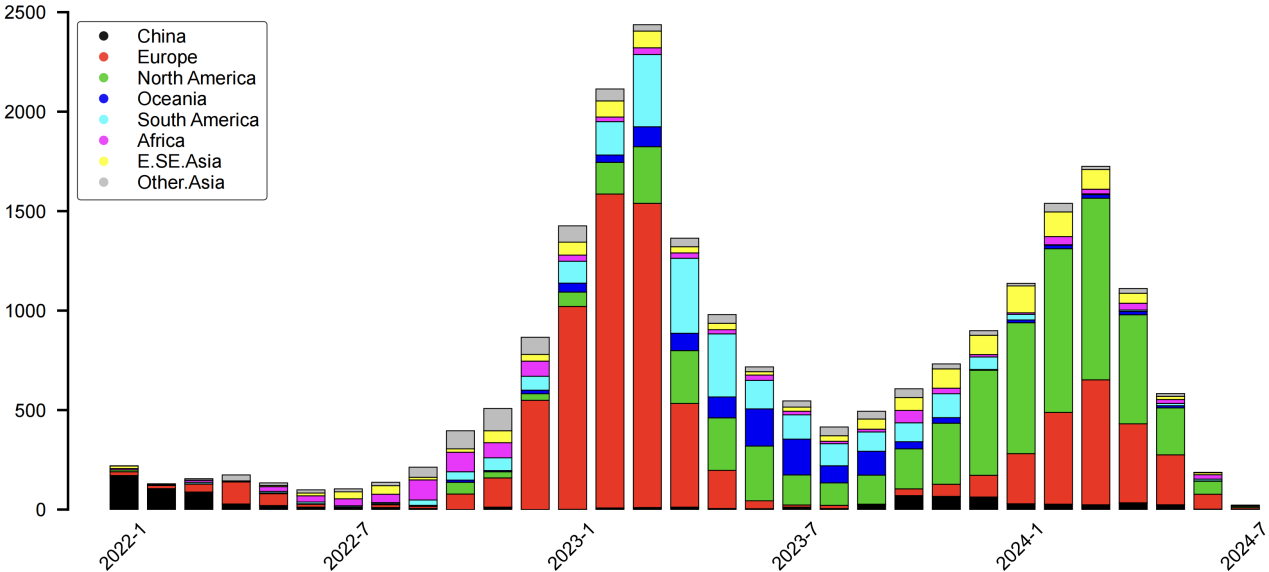

**Figure S6**. The phylogenetic tree represents recent influenza B (Victoria clade) viruses, with colors indicating the clade (left figure) or antigenic clusters (middle figure) as per the legend in the top-left corner of the figure. The right figure refers to the isolation time of viruses in the phylogenetic tree. Q1 and Q2 refer to the first and second quarter of this year. Clades were determined using the Nextclade pipeline.

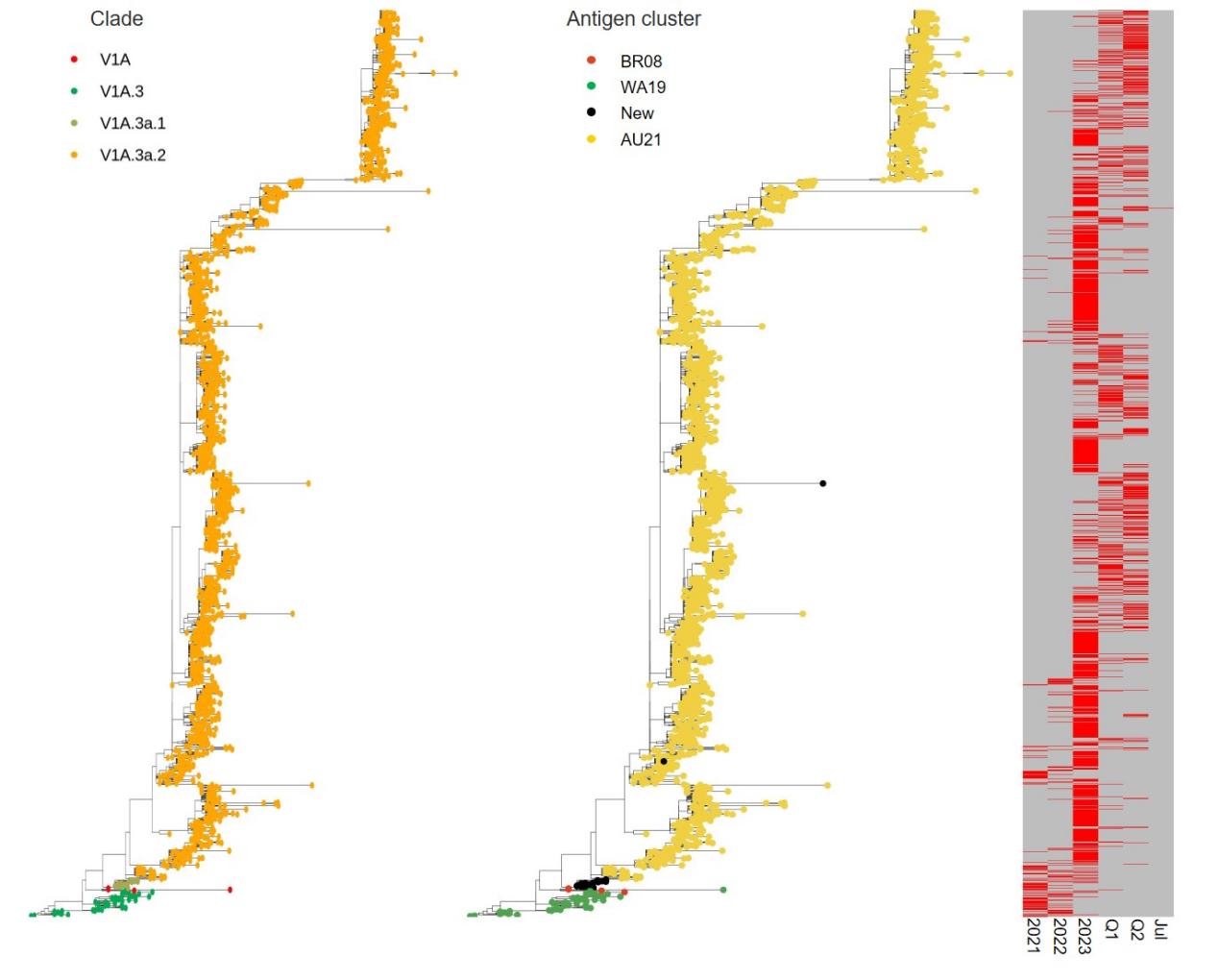

**Figure S7**. The monthly ratio of antigenic similar viruses to the latest vaccine strain B/Austria/1359417/2021 in the last two years.

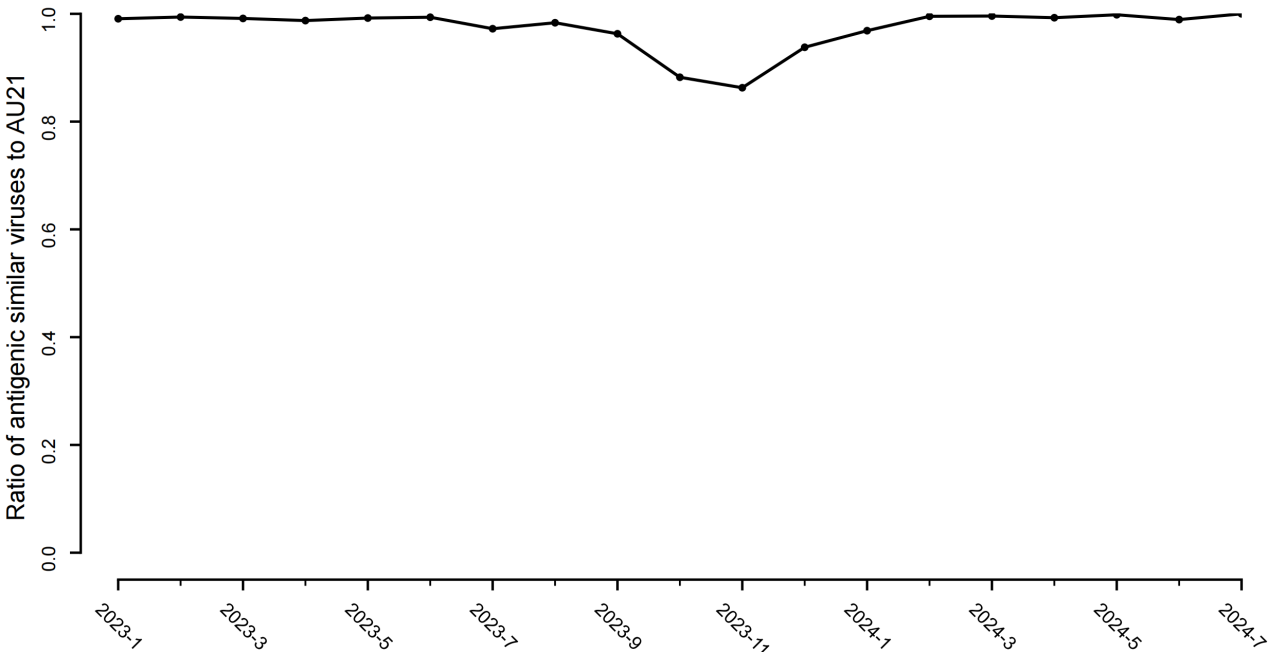
